# Investigating the demographic history of Sindhi population inhabited in West coast India

**DOI:** 10.1101/2025.03.01.640946

**Authors:** Lomous Kumar, Suraj Nongmaithem, Sachin Kumar, Kumarasamy Thangaraj

## Abstract

**Background:** South Asian populations are genetically well stratified due to multiple waves of migration, admixture events, and endogamy. India remains a rich resource for population genomics studies with many small and socioculturally homogeneous communities whose origins and demographic histories are largely unknown. In this study, we analysed such a small Sindhi settlement in the Thane district in Maharashtra of West coast India using genome-wide autosomal SNP data using both frequency- and haplotype-based approaches.

**Results:** Our analyses suggest that the West coast Indian Sindhi community is very unique and has significant population affinity with a group more closely related to the Pakistani Burusho than to the Pakistani Sindhi, as it has an additional East/Southeast Asian component. Furthermore, the sharing of haplotype and IBD suggests recent gene flow from the local Konkani population on the west coast of India into Indian Sindhi. Admixture modelling suggested that Indian Sindhi admixture with the East/Southeast Asian source group could be 40-50 GBP, explaining their current unique demographics. However, apart from this additional admixture, they share the basic genetic composition of the Pakistan/NWI groups, as reflected in PCA, outgroup F3 and IBD sharing.

**Conclusion:** These new findings suggest that Indian Sindhi settlement from the Thane in Maharashtra in West coast of India derive their genetic ancestry not directly from Pakistani Sindhis but from other groups related to Burusho in Pakistan. The study therefore encourages further research to identify the heterogeneous nature of migrations to the Indian subcontinent and thus further decipher its unique demographics.

## Introduction

South Asia is considered as one of the most geographically diverse locations in terms of linguistics, culture, and genetics (1, 2). Among the South Asian countries, India and Pakistan are the land of fascinating diversity, a vibrant web of different cultures, languages, traditions and beliefs (3). Archaeological finds from the Mesolithic and Palaeolithic periods in India and Pakistan represent crucial phases in human prehistory characterized by significant migrations, technological advances, and cultural developments (4-6). The archaeological evidence of various sites, i.e. H. Paleolithic sites such as Soan Valley (Pakistan), Bhimbetka (India), etc., and Mesolithic sites such as Bagor (India), Mehrgarh (Pakistan), etc., across the region provide valuable insights into the lives and movements of early human populations (1, 7, 8). The region encompassing present-day India and Pakistan is home to some of the oldest and most advanced civilizations in human history, such as the Indus Valley Civilization (IVC - c. 3300-1300 BC), the Vedic Civilization (c. 1500-500 BC) and the Mauryan Empire (ca. 322–185 BC), Gandhara civilization (ca. 1500 BC–500 AD) (9-14). The remarkable diversity of the population in this region is evidence of its rich history, centuries of cultural exchange and coexistence of numerous ethnic groups (2, 15) These geographical strata (i.e. India and Pakistan) play an important role in the distribution of human migration, which leads to the exchange of culture and language according to different ethnic populations (5, 16).

One of the largest, fastest and most recent human migrations was observed during the partition of India in 1947 (17, 18). This mass movement of people was not only a population shift but also an ethnic and religious reconfiguration of the region, with millions forced to leave their ancestral homes (19, 20). Among the various affected populations, the Sindhis living in Pakistan’s Sindh province were particularly affected (21). Sindh province utilizes one of the ancient archaeological sites (3rd millennium BC) from the Indus Valley Civilization (IVC), namely Mohenjo-Daro (22). During Partition, a large number (about a million) of non-Muslim Sindhi migrated to Gujarat, Rajasthan, and Punjab (23, 24). They speak the Sindhi language, which belongs to the Indo-European language family. Over time, the language underwent various transitions such as Prakrit, Arabic, Persian, etc. due to the influence of different traders and rulers (25, 26). From its ancient roots, such as the Indus Valley Civilization, to its present status, Sindhi has absorbed and integrated influences from various languages and cultures, making it a unique and resilient language (27).

The genetic makeup of the Sindhi population also reflects historical migrations and interactions with neighbouring regions (28). Genetic studies on the maternal markers of the Sindhi population show a wide range of mtDNA haplogroups, indicating diverse maternal ancestry (29). Many of the mtDNA haplogroups in Sindhis are ancient and have deep roots in the South Asian subcontinent, such as M, R and U (30). In contrast, haplogroups such as W and HV indicate genetic links with populations from West and Central Asia (29, 31). Genetic markers shared with North Indian (Indo-European) and South Indian (Dravidian) populations indicate a history of gene flow between these groups in Sindhis from South Asia (32). The Arab conquest of Sindh in the 8th century introduced new maternal lineages into the Sindhi gene pool, reflected in the presence of West Eurasian haplogroups (29, 31, 33).

The paternal ancestry of the Sindhi population, like their maternal ancestry, reflects a rich and diverse history marked by ancient migrations, invasions and cultural interactions (34, 35). The common haplogroups among Sindhis include R1a, J2, J1, L and H, which are widespread in South Asia and adjacent regions (35-38). R1a in Sindhis indicates the influence of Indo-European migrations into the subcontinent about 3,500 years ago, J2 is associated with agricultural communities in the Fertile Crescent and ancient Mesopotamia. Haplogroup L is widespread in the Indian subcontinent and the Middle East (36, 39). This haplogroup is widespread in South Asia and is associated with the Dravidian populations; The presence of J1 haplogroups is typical of the Arabian Peninsula, which is less common but is still notable among Sindhis (37, 39).

The studies on autosomal STR markers also suggest a connection between Sindhis and other populations of Pakistan and Northwest India (38, 40). Other genome-wide studies have also shown a relationship between northwest Indian populations such as Gujjar and Ror, and the Sindhis population, proving that they share genetic ancestry with Indo-Europeans and Middle Eastern populations (41). These connections likely stem from trade, invasions, and other forms of interaction over millennia. Earlier study pointed out the connection between Northwest India (NWI) and Southwest coast India (42) through much earlier migrations. However, very limited genetic information available about Sindhis living in other parts of India, particularly in the West coast part of India. In the present study, for the first-time, we report genotype data of the Sindhi settlement from the Thane in Maharashtra in the West coastal region of India. This study will examine the common ancestry, assimilation and the past migration history of the Sindhis to South India.

## Methods

For this project, we followed approved guidelines, applied protocols, and obtained approval from the Institutional Ethics Committee of CSIR-CCMB, Hyderabad, India. Blood samples were collected from 13 healthy Sindhi individuals (all Male) from Thane in Maharashtra, India after written informed consent was obtained and DNA was isolated using the standard phenol-chloroform method and used for subsequent genetic analysis. All Indian Sindhi samples (n=13) were genotyped with the Illumina HumanOmniExpress 24 v2 array kit using the manufacturer’s protocol for a total of 642,824 genome-wide single nucleotide polymorphisms. The dataset was merged with the published DNA dataset of contemporary populations from the HGDP(43) and Genome Asia panel (44) and quality filtering was applied in Plink 1.9 (45) to include only autosomal markers on 22 chromosomes with a genotyping call -Rate of > 99% to be retained and minor allele frequency >1% (511668 SNPs). Kinship based filtering was performed by removing individuals with first- and second-degree relatives using the KING-robust function implemented in Plink2 (45, 46). After all filtering, the final merged dataset included 866 modern individuals genotyped at 511,828 SNPs. To minimize the effect of background LD in PCA and ADMIXTURE-like analyses (47), markers were further pruned by selecting SNPs in strong LD (r2 > 0.4, window of 200 SNPs, sliding window of 25 SNPs each) using Plink 1.9 (45). The LD-trimmed data included 365,621 SNPs on the autosome. For all subsequent analysis except PCA and admixture, the full SNP set of 511,828 sites was used.

Principal component analysis (PCA) was performed on the merged dataset of modern Eurasia using the Smartpca package implemented in EIGENSOFT 7.2.1 (48) with default settings. The first two components were recorded to infer genetic variability. The model-based clustering algorithm ADMIXTURE (47) was executed to infer ancestral genomic components in the Indian Sindhi population. Cross-validation was performed 25 times for 11 ancestral clusters (K=2 to K=12) (Fig. S1). The lowest CV error parameter was obtained at K = 6 and used for downstream analysis. The qp3Pop implementation of the ADMIXTOOLS package (49) was used to calculate the outgroup F3 statistics. To infer the gene flow of modern Eurasians in the Indian Sindhi population, the F3 statistic was used in the form F3 (Mbuti; SND, X), where X is any modern West Eurasian or South Asian population. (SND = Indian Sindhi).

The haplotype-based approach implemented in CHROMOPAINTER (50) and FineStructure (50) was used to infer a fine-scale co-ancestry matrix and population clustering, respectively.

The data were first phased with SHAPEIT5 (51) using default parameters, followed by a CHROMOPAINTER run to derive the co-ancestry matrix, first by performing a 10-expectation maximization iteration (EM) with 5 randomly selected chromosomes with a subset of individuals to derive global mutation rate (µ) and switch rate (Ne) parameters. The main algorithm was then run on 22 chromosomes from all individuals to derive the co-ancestry matrix. This matrix was used by FineStructure to infer clustering using a probabilistic model by applying the Markov Chain Monte Carlo (MCMC) method and then deriving a hierarchical tree by merging all clusters with the least change in posterior probability. The run used 500,000 burn-in iterations and 5,000,000 subsequent iterations and the results of each 10,000-iteration saved. Estimates of admixture date and best admixture models were derived with fastGlobeTrotter (52) using Chromopainter chunklength files.

For identity by descent (IBD) analysis, we performed haplotype inference or phasing using three independent runs of Beagle-5.4 (53). IBD segments were determined from phase data from all three runs separately using refined IBD (54), then segments from all three runs were combined, and then combined segments were merged using the Merge IBD Segments tool. The IBD release matrix was then recorded using a custom script in R (55).

To derive the best-fitting demographic model and model parameters, we used the parameter optimization method implemented in Moments (56). For Indian Sindhi, we used a preliminary model based on hypothesis driven from either FastGlobeTrotter (52) admixture models of Indian Sindhi groups, with much earlier Dai-like admixture or alternatively a more recent pulse in India. For model construction, we used the Python package Demes (57). Parameter files were created based on the respective Demes models. Two alternative models were used to compare the demographic scenario of Indian Sindhi groups (Supplementary Fig S4 & S5). The site frequency spectrum was calculated from empirical data in VCF format as well as from Demes model specifications using moments. Model parameter optimizations were performed with 500 iterations and the *lbfgsb* method. Confidence intervals for derived parameters were calculated using the *moments*.*Demes*.*Inference*.*uncerts* function of Moments (56).

## Results

### Genetic structure in Indian Sindhi

The PCA biplots presented includes Indian Sindhi (black), Indo-Europeans (blue), Dravidians (red), Austroasiatic speakers (khaki), Tibeto-Burman (orange), Pakistani groups (forest-green), and North West Indians (light-green) (Fig 1b). Interestingly, the Indian Sindhi (black dots) clustered near one extreme of the South Asian cline (but away from main cline) with most of the Pakistan and Northwest Indian (NWI) groups with highest ANI ancestry. Their clustering pattern was more shifted towards the Burusho from Pakistan. This shifting is towards the PCA axis occupied by East/Southeast Asian groups (Fig 1b). Two of the Indian Sindhi individuals are shifted towards Indian Indo-Europeans and one of them clustered along with individuals from Konkani population.

**Fig 1:**
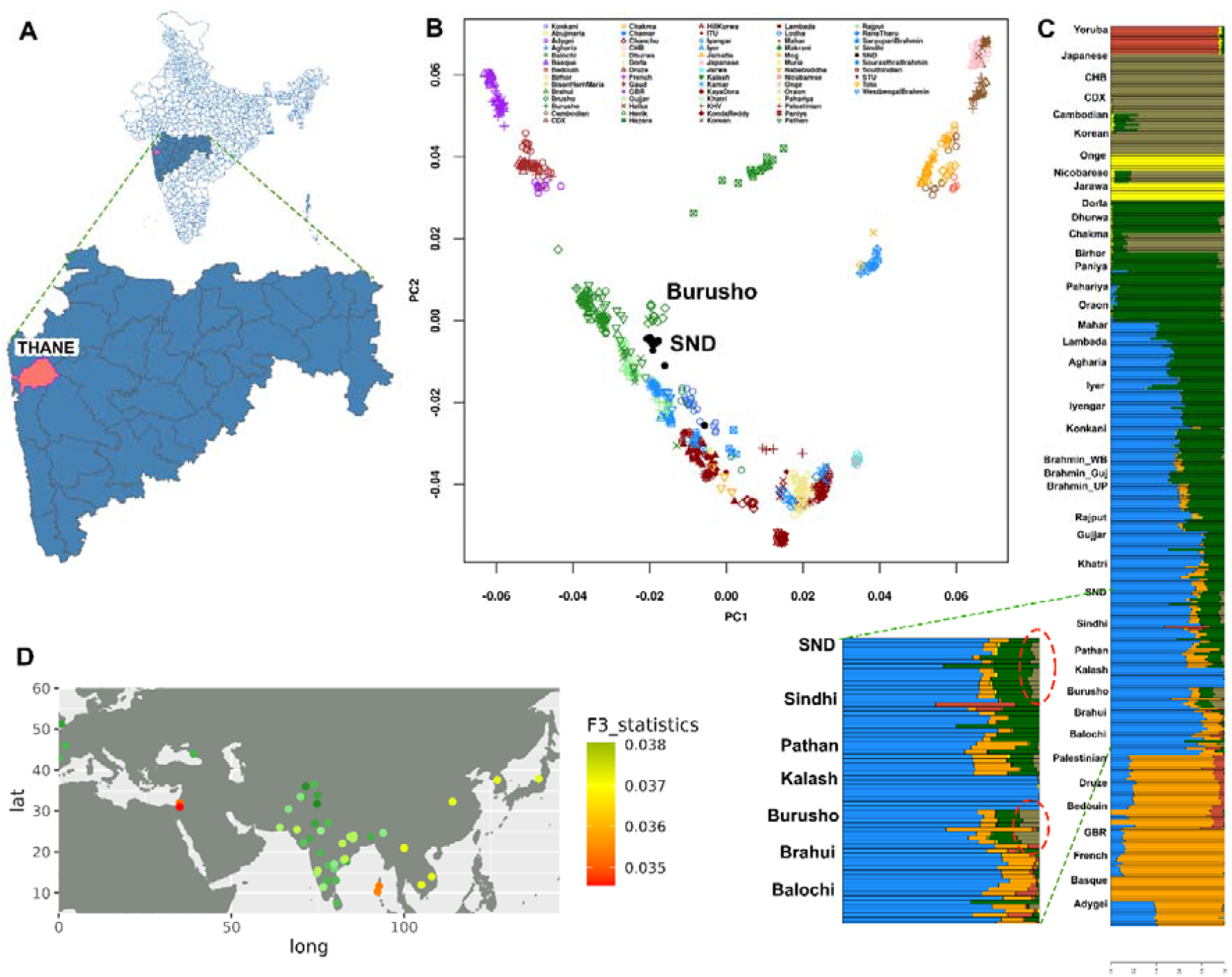
Sampling location and population structure of Indian Sindhi. **A**. Location in the Indian state of Maharashtra (Thane) from West coast, **B**. PCA biplot of Indian Sindhi (SND) with modern Eurasians, **C**. Admixture barplot with modern Eurasians (red elipse shows East Asian Khaki colour component). **D**. Outgroup F3 statistics with modern Eurasians (showing highest allele sharing of SND with NWI) (Dark green colour in NWI).

In the unsupervised model-based clustering with ADMIXTURE (47) using K=6 (Fig. 1C), Indian Sindhi formed a unique East Asian component (Khaki) different from most of the Pakistan/NWI groups but this component was also observed in Burusho and only few individuals of Pathan. This ancestral component is maximised among East Asians, Southeast Asians, Indian Tibeto-Burmans and Indian Austroasiatic groups. Besides this both Indian Sindhi and Burusho have typical South Asian Indo-European ancestral components (Blue, Forest green and Orange) (Fig. 1C).

### Allele sharing between Indian Sindhi and modern Eurasians

In the outgroup F3 statistics, the Indian Sindhi showed highest allele sharing with the Khatri population (F3 = 0.2648; z = 133.8107) from NWI followed by Kalash (F3 = 0.02639; z = 129.6765) from Pakistan and Gujjar (F3 = 0.02621; z = 133.5288) again from NWI (Supplementary_Table S1). Some of the other top hits were mostly Indian Indo-European populations like Rabadi, Rajput and Brahmin_UP.

### Fine scale population structure and IBD sharing

The fineSTRUCTURE (50) tree kept all the populations in two major clades, with one major clade included East/Southeast Asians, Andamanese, Indian Tibeto-Burman, Indian Austroasiatic and Dravidians, while other clade incorporated Europe/MidEast, Pakistan/NWI and Indian Indo-Europeans (Fig 2a). In this second clade Indian Sindhi forms an altogether separate minor branching with Konkani group from west coast India. Most of the Pakistani Sindhi individual were sharing clades with Pakistan/NWI populations (Fig 2a).

**Fig 2:**
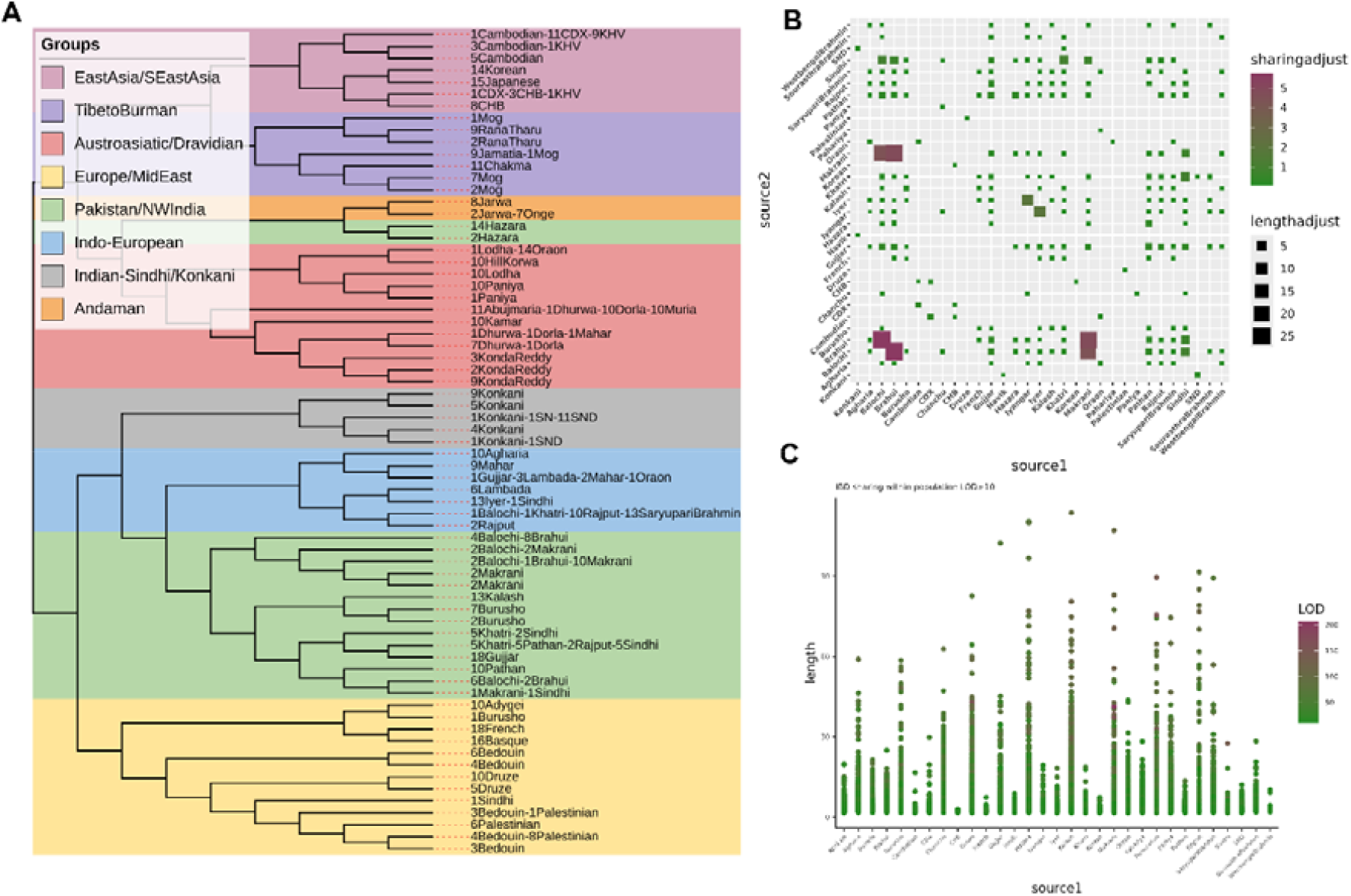
Haplotype and IBD sharing statistics. **A**. fineStructure MCMC tree for Indian Sindhi (SND) with all modern Eurasians, **B**. Inter-population IBD sharing matrix adjusted for population size, **C**. Intra-population IBD sharing within group (LOD > 10).

In the cross population IBD sharing matrix adjusted for population size, Indian Sindhi showed highest IBD sharing with Konkani population from West coast (Maharashtra) India, followed by Khatri population from Northwest India (NWI) (Fig 2b). Whereas, in the intra-population IBD sharing the length distribution is smaller in comparison to most of the modern Eurasians and almost comparable to Pakistani Sindhi (Fig 2c). In the within population IBD sharing, we excluded the values below LOD score of 10 in all cases.

### Admixture modelling and dating

The best fit sources for Indian Sindhi in the estimation of the best fitted model and date of admixture using fastGlobeTrotter (52) were Khatri from Northwest India (NWI) and Dhurwa (an Austronesian proxy) (Fig 3b). The best fit date of admixture was approximately 46 GBP (Supplementary Table S2).

**Fig 3:**
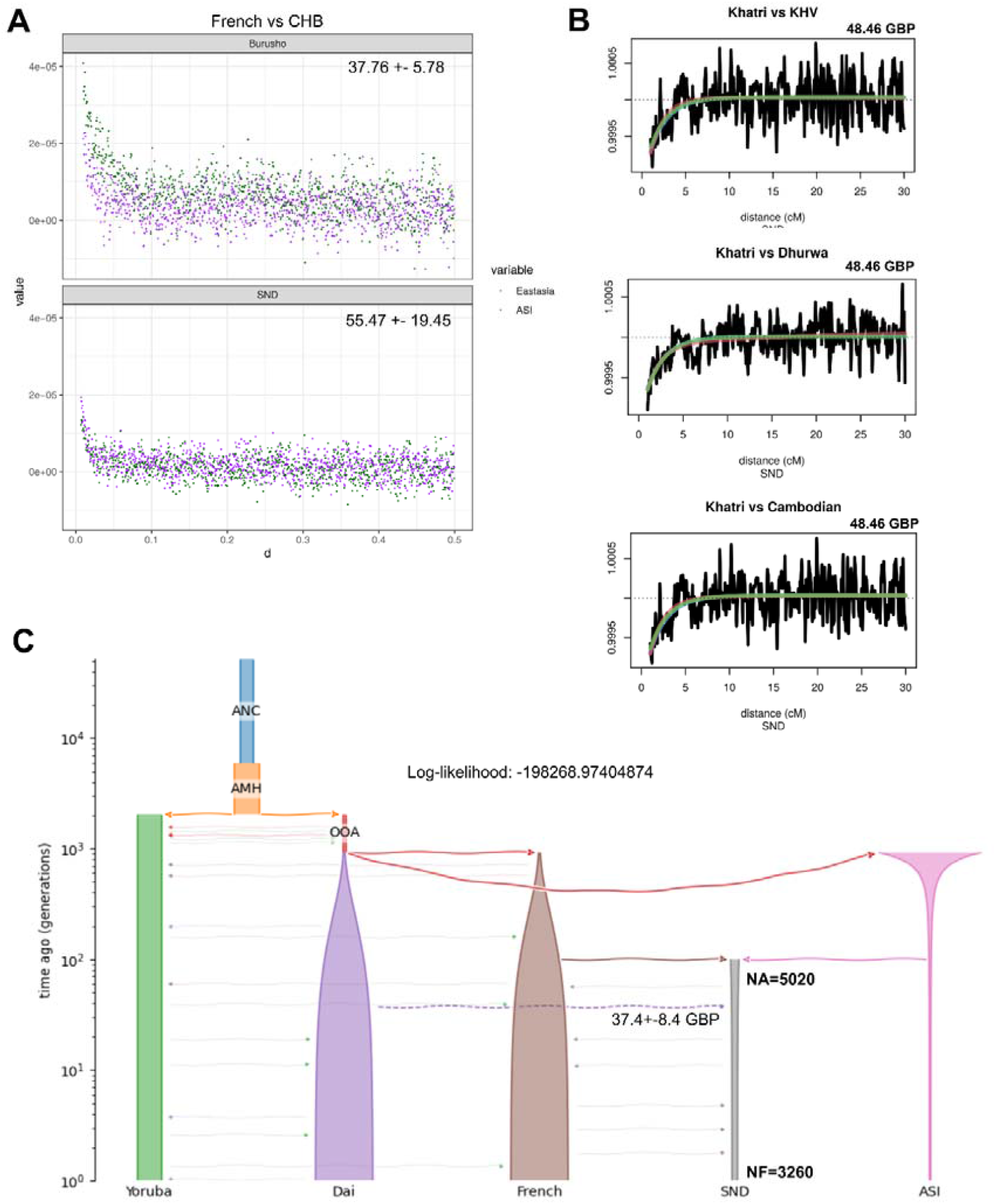
Admixture modelling and dates of Indian Sindhi **A**. ALDER LD decay curve for Burusho and SND (Indian Sindhi), **B**. fastGlobeTrotter co-ancestry curve fitting for SND (Indian Sindhi), **C**. Best fit demographic model with admixture date from Dai in SND (Indian Sindhi).

### Demographic history and demographic parameter estimation

We proposed two alternate demographic models for model competition for the Indian Sindhi population, with first model based on our admixture modelling results of Dai admixture, which is evident in model-based Admixture as well as fastGlobeTrotter (52) modelling and another model with possible admixture with Indian (possibly Austroasiatic) groups as source of Dai-like component. For replicating the admixture history of Indian Sindhi in the prior model, we used Demes (57). We used Moments’(56) *inference optimization* function to arrive at best likelihood model and parameters. Of the tested two alternate models, Model1 hypothesize the pulse of admixture from Dai-like source much earlier and with similar event to Burusho in Pakistan (probably through Mongolian invasion), while Model2 hypothesize putative admixture between ANI-ASI and later admixture event with Dai-like source recently from Indian Austroasiatic groups to form the Indian Sindhi group. We selected Model1 (Log-likelihood: -198268.97404874387) (Fig 3c) over Model2 (Log-likelihood: -206243.77405798397) based on their likelihood scores, which corroborated well with admixture model inferred from fastGlobeTrotter (52) run. This recent admixture with Dai was dated to approximately 37.4 GBP (95% C.I. 29.003-45.8005), which is almost comparable to the date estimate in Burusho population from fastGlobeTrotter run and upper limit of 95% C.I. corresponds to Indian Sindhi fastGlobeTrotter estimate (46 GBP) (Supplementary Table 1b).

The parameter estimates from best fitted model of Indian Sindhi suggest that there was not significant reduction in effective population size in this group (NA=5200; NF=3260), with noticeable migration rates between ANI and Indian Sindhi (M_French_GroupA =0.00183) (Fig. S3). This effective population size change was more prominent in case of ASI (Paniya as a proxy; Ne=61000 and NeF=854) (Fig 3c).

## Discussion

The high level of population diversity and stratification in India is due to multiple waves of migration from outside into the region over millennia and eventual mixing and cultural assimilation. Genetic evidence of later migrations and admixture events is well documented, particularly on the west coast of India, such as among Indian Parsees (Chaubey, Ayub et al. 2017), Cochin Jews (Chaubey, Singh et al. 2016), and Roman Catholic Jews (Kumar, Farias et al. 2021) and migration and local assimilation of warriors clans from Northwest India to Southwest coast (42). In the present study, we have carried out, for the first time, a detailed investigation of the genetic architecture of a small, isolated and socio-culturally unique Indian Sindhi settlement from the Thane in Maharashtra. Our analysis suggests that This Indian Sindhi settlement in Thane represent a unique group, distinct from the local population in India as well as the Pakistani Sindhi population. Their genetic structure surprisingly indicates their closer affinity to the Burusho-like population from Pakistan, due to the presence of an additional East/Southeast Asian genetic component. This Indian Sindhi subgroup did not form a close group with the Pakistani Sindhi, reflecting their marked contrast with that group. Allele sharing statistics (outgroup F3) using Mbuti as the outgroup and various modern Eurasians as the test group indicate greater shared-drift of Indian Sindhi with populations from Northwest India (NWI) and Pakistan. This reflects their long-standing shared affinity with Pakistan/NWI, as does Pakistani Sindhi. This pattern of genetic affinity of populations from Pakistan (Sindhi, Pathan etc.,) with Northwest Indian is well pointed out through earlier genetic studies (38, 40, 41). Indian Sindhi have a well-documented migration history from Pakistan/NWI, which correlates well with our genetic findings.

Of note, the genetic architecture of these Indian Sindhis from Thane in context of local population also equally draw attention, as they share haplotypes with the local Konkani groups from West coast India. This kind of haplotype sharing often reflects very recent gene flow patterns and this is also evident in the IBD chunk sharing. They share larger IBD segments with Konkani population apart from comparatively short segments with Khatri population from Northwest India (NWI). Former represents much recent gene flow while later represents earlier shared genetic history of Indian Sindhi with NWI populations. Pakistani Sindhi also share their genetic history with Pakistan/NWI populations based on earlier study on Y-STR (Anwar et al. 2019; Perveen et al. 2017). Further Northwest Indian populations like Ror and Gujjar showed major affinity with Pakistan and Northwest Indian populations (41). Therefore, the PCA-based clustering of Indian Sindhi among Pakistani populations as well as haplotype sharing with Khatri largely reflects their long-term shared genetic history with both Pakistani and northwest Indian populations.

At this point, it is important to discuss the unique East Asian (Dai-like) component that clearly distinguishes Indian Sindhi settlement in Thane from Pakistani Sindhi. Our haplotype-based admixture modelling suggests that this minor component (∼10%) was introduced at 46.46 GBP with the best-matched surrogates as Khatri (NWI) and Dhurwa (Austroasiatic group). The latter group represents a proxy for an Austronesian (Dai-like) surrogate. Estimates of admixture date using the linkage disequilibrium-based method were also similar (55.47+-19.46). Furthermore, these date estimates were well supported in our demographic modelling, with the admixture timing found to be 37.4 GBP from the Dai-like source group in the best-fit model. Second model which suggest more recent gene flow from a group carrying Dai-like component after Indian Sindhi migration to India, is excluded. Thus, successful model indicates much earlier admixture of this component and the time frame overlaps with the Mongolian (Genghis khan) invasion in Pakistan although other possible source cannot be excluded. Apart from Indian Sindhi and Burusho, Pathan also shows the minor presence of this additional East Asian component. The Late Bronze Age Steppe populations and many Iron age migrations (Saka, Hun, Kushan etc) were having additional East Asian genetic components (58). Most of the Populations from Pakistan (Balochi, Pathan, Burusho, Sindhi and Hazara) are in geographical proximity and in a transition zone in relation to pre-Historical and Historical migrations. Hence there is possibility of acquiring such component during any of these migration waves, which require much detailed genetic investigations. Although the ALDER-based estimate was comparatively at the high end, the fastGlobeTrotter date estimate was at the upper limit of the 95% confidence interval of the demographic model-based admixture date estimate (95 % C.I. 19-45 GBP). This may be due to other admixture events or noise in ALDER admixture date estimates. Furthermore, parameter estimates in our Indian Sindhi demographic modelling revealed no evidence of a significant founding event or population bottleneck in this group, and there was no significant change in the effective population size.

In conclusion, this study presents the first insightful genetic evidence for the ancient origin and unique genetic architecture of Sindhi population settlement from the Thane in Maharashtra in West coast India. The study is the first to report evidence of East Asian admixture in this group much earlier in history than their migration to the West coast of India. Their genetic assimilation with the local majority population (Konkani) reflects their predominantly exogamous nature, which is well reflected in lower IBD exchange within the population. Given the limited sample size of the Indian Sindhi from West coast India in the present study, future efforts with a much larger sample size incorporating Sindhi populations from most of the India along with uniparental and whole genome analysis will reveal more interesting aspects of their population history. Furthermore, further population genetic studies of this kind on different groups will shed light on the heterogeneity of many similar genetic migrations to India.

## Ethical Approval

Informed written consent was obtained from each participant. The project was carried out in accordance with the guidelines approved by the Institutional Ethical Committees of Centre for Cellular and Molecular Biology, Hyderabad, India. All the procedure has been followed according to the recommendations of the Helsinki Declaration.

## Declaration of interest

The authors declare no competing interests.

## Informed consent

Informed written consent was obtained from all the participants involved in the study.

## Data availability Statement

The data supporting the findings of this study are available upon request.

## Acknowledgements

We thank all the study participants, who volunteered in this study. KT was supported by J C Bose Fellowship (JCB/2019/000027) from the Science and Engineering Research Board (SERB), Department of Science and Technology, Government of India.

## Author contributions

KT conceptualised the study and recruited the study samples. LK and KT devised the methodology. LK and SN genotyped the genome wide SNP markers. LK performed the data analyses. KT and LK wrote the first draft of the manuscript. KT finalised the report. KT provided feedback on the report. All authors contributed to and have approved the final manuscript.

## Notes

### Competing Interest Statement

The authors have declared no competing interest.

