## Supplementary Figures for "Investigating the demographic history of Sindhi population inhabited in West coast India"

**Supplementary Fig1**: Biplot of first two principal components in the PCA analysis of Indian Sindhi with all modern Eurasians including African populations. (**Indian Sindhi/SND represented by black colour points**)

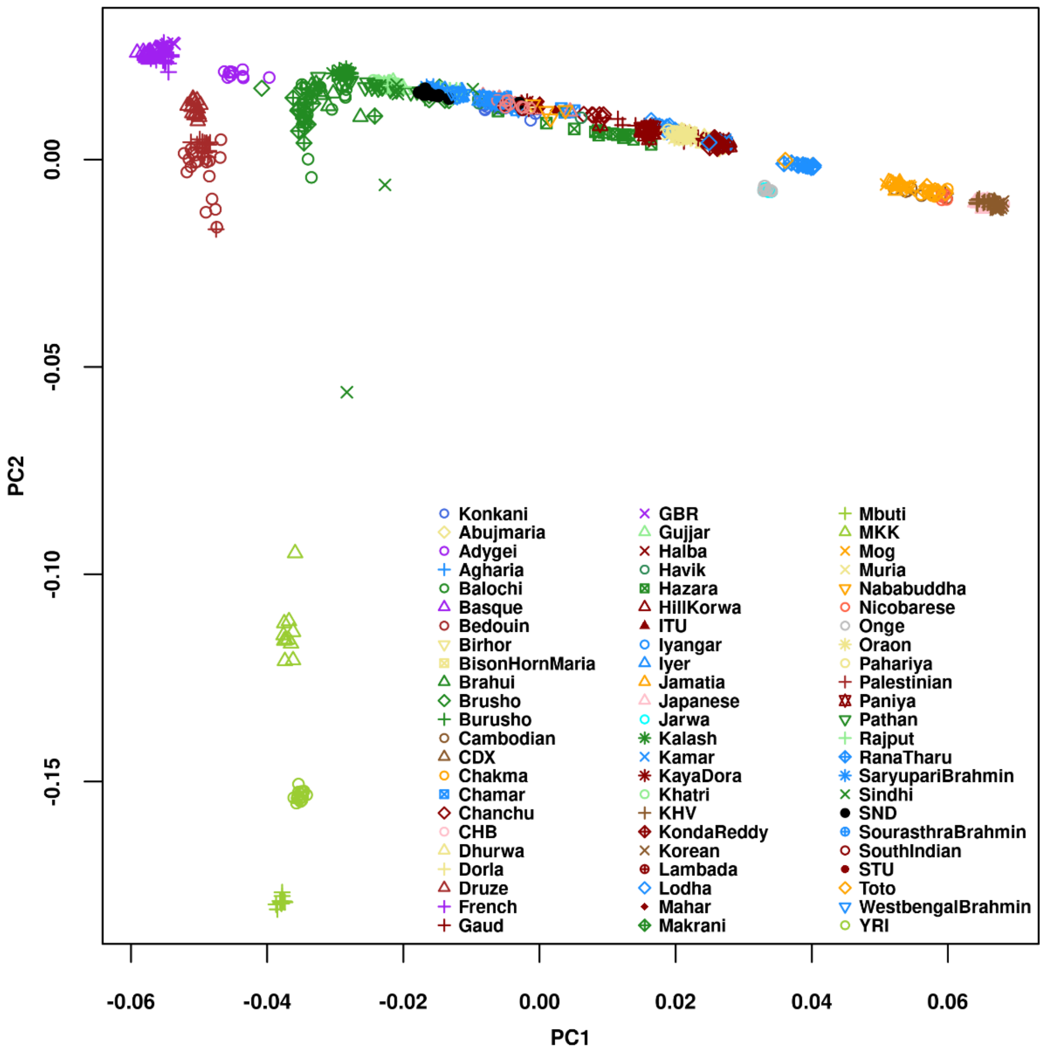

**Supplementary Fig 2**: CV error plot of cross validation run of model-based clustering using Admixture. Plot shows the lowest cv error at K value of 6.

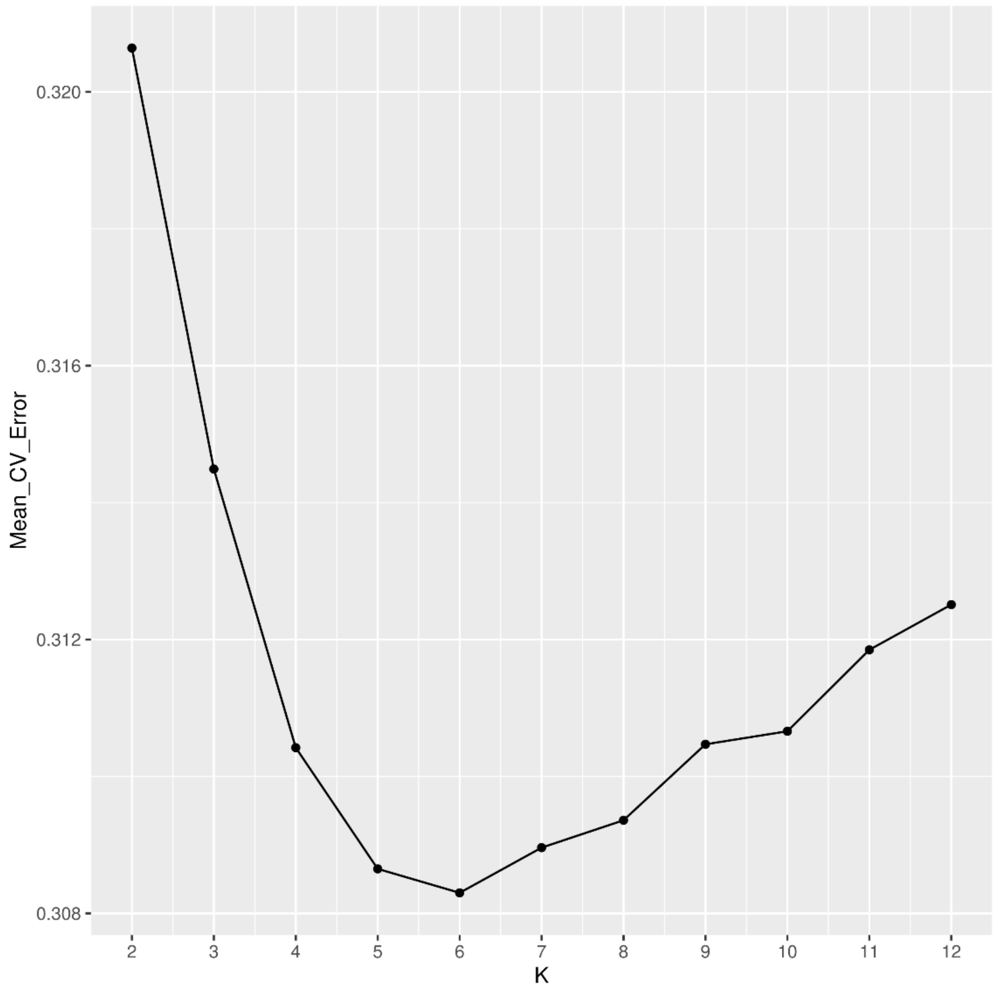

**Supplementary Fig 3**: Co-ancestry matrix of Chromopainter chunk sharing of modern Eurasians with Indian Sindhi (**SND**).

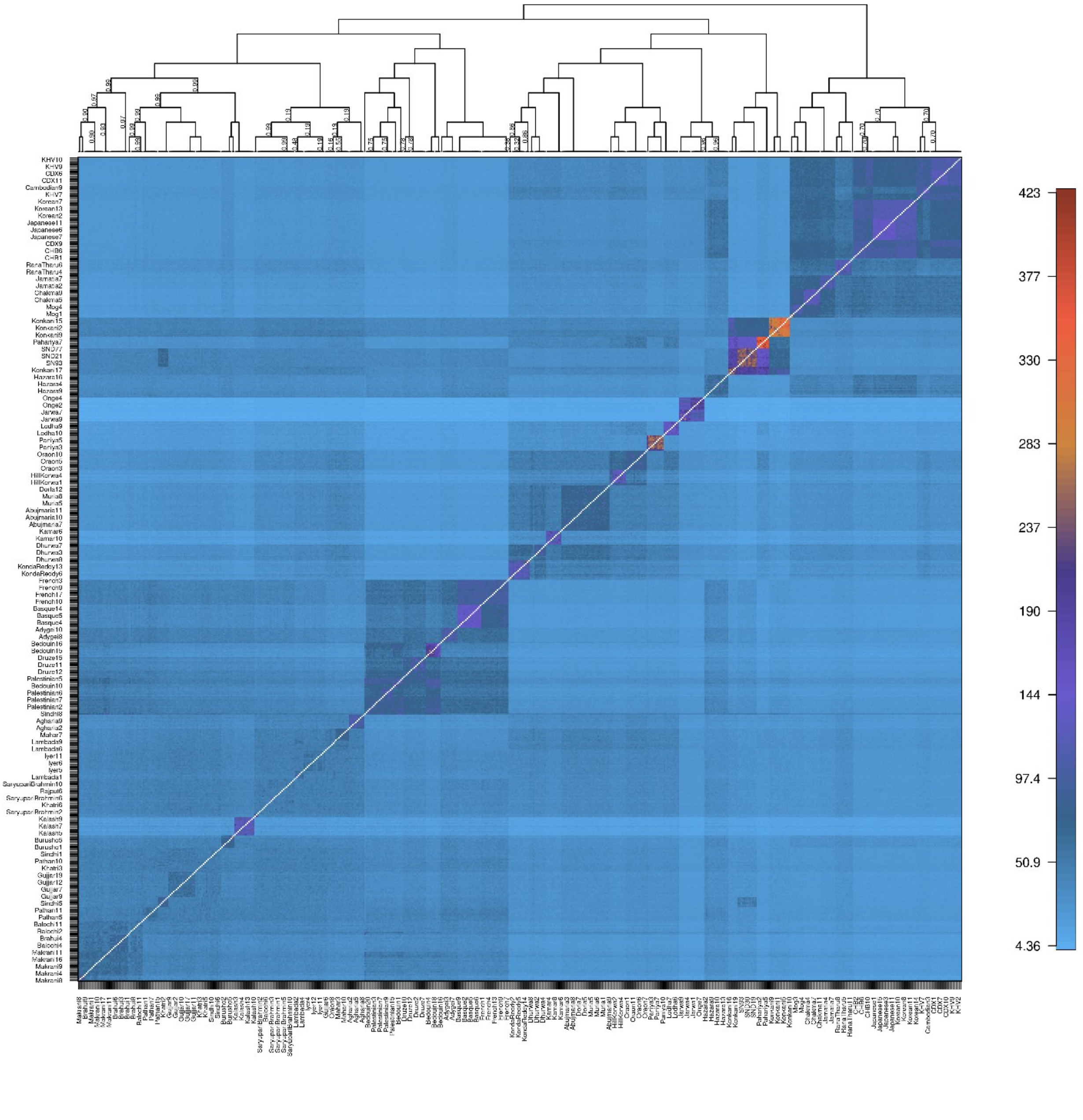

**Supplementary Fig 4**: Demes-based preliminary model1 used for the fitting the demography of Indian Sindhi (SND). (Dai related ancestry is shown to be introduced much earlier and much similarly like Burusho population).

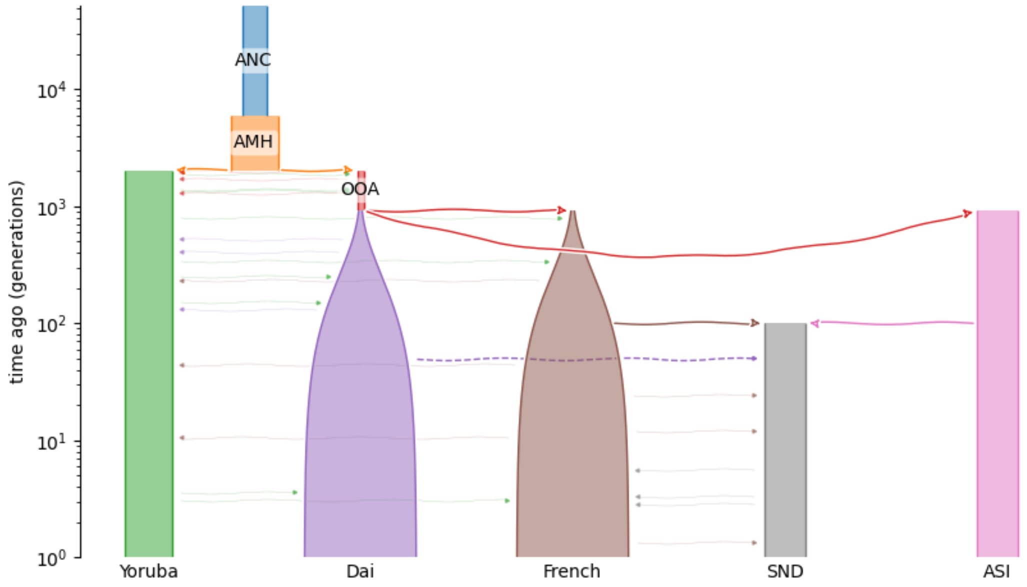


**Supplementary Fig 5**: Demes-based alternate preliminary model2 tested to infer the demography of Indian Sindhi. (Dai related admixture is shown to very recently introduced via Indian Austronesian ancestry source or Austroasiatic source).

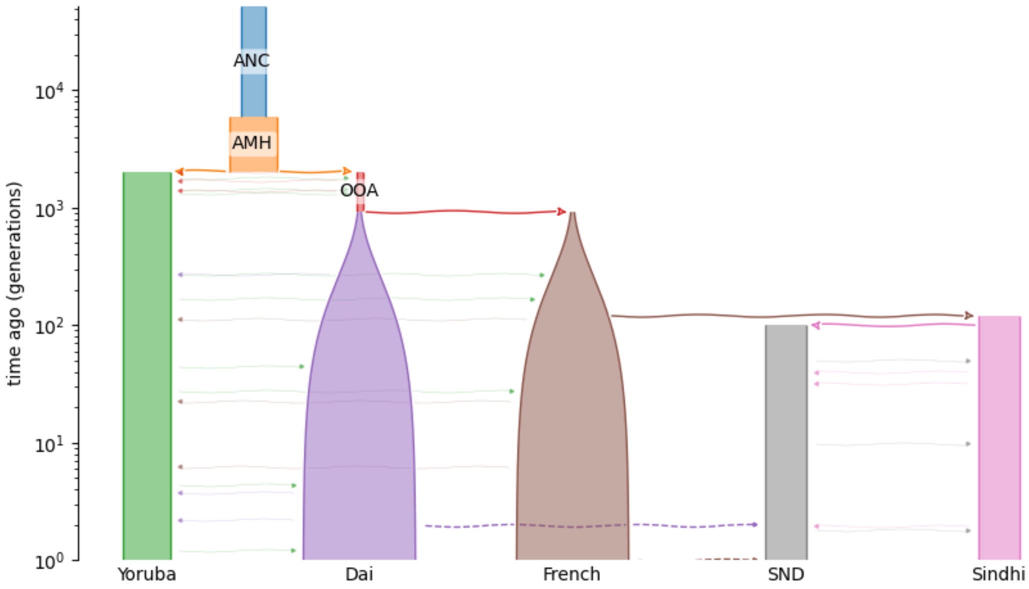

**The model fits and parameters for two alternate models tested for East/Southeast Asian pulse in Indian Sindhi:**

##### Model1 ######

Log-likelihood: -198268.97404874387

Best fit parameters

Ne 6.1e+04 (Initial effective population size of ASI)

NeF 8.54e+02 (Final effective population size of ASI)

NA 5.02e+03 (Initial effective population size of Indian Sindhi)

NF 3.26e+03 (Final effective population size of Indian Sindhi)

M_French_SND 0.00183 (migration rate between ANI and Indian Sindhi)

Tadmix_Dai_SND **37.4** (pulse of admixture from Dai to Indian Sindhi)

95% CIs

| param | 2.5% | 97.5% |
|---|---|---|
| Ne | 58260.3 | 63755.5 |
| NeF | 840.289 | 868.362 |
| NA | nan | nan |
| NF | nan | nan |
| M_French_SND | 0.00174679 | 0.00191117 |
| Tadmix_Dai_SND | **29.003** | **45.8005** |

###########Model2 ########

Log-likelihood: -206243.77405798397

Best fit parameters

Ne 4.12e+03 (Initial effective population size of ASI)

NeF 3.91e+03 (Final effective population size of ASI)

NA 2.95e+03 (Initial effective population size of Indian Sindhi)

NF 2.23e+03 (Final effective population size of Indian Sindhi)

M_Sindhi_SND 0.0349 (migration rate between Pakistan SIndhi and Indian Sindhi)

Tadmix_Dai_SND 6.82 (pulse of admixture from Dai to Indian Sindhi)

95% CIs

param 2.5% 97.5%

| | | |
|---|---|---|
| Ne | 4110.51 | 4127.02 |
| NeF | 3875.24 | 3940.88 |
| NA | 2929.73 | 2962.09 |
| NF | 2199.55 | 2269.44 |
| M_Sindhi_SND | 0.0348298 | 0.0350492 |
| Tadmix_Dai_SND | 6.7118 | 6.9197 |
